# Quantifying brain-wide cerebrospinal fluid flow dynamics using slow-flow-sensitized phase-contrast MRI

**DOI:** 10.1101/2025.03.22.644745

**Authors:** Zijing Dong, Fuyixue Wang, Amelia K. Strom, Korbinian Eckstein, Beata Bachrata, Simon D. Robinson, Bruce R. Rosen, Lawrence L. Wald, Laura D. Lewis, Jonathan R. Polimeni

## Abstract

Cerebrospinal fluid (CSF) flow is a key component of the brain’s waste clearance system. However, our understanding of CSF flow in the human brain, particularly within the brain-wide subarachnoid space (SAS), is limited due to a lack of non-invasive tools for measuring slow flow. Here, we propose a CSF flowmetry technique using phase-contrast MRI combined with a slow-flow-sensitized acquisition. It achieves high sensitivity in measuring slow CSF flow (e.g., 100 μm/s), and enables quantitative measurement of the velocity and direction with whole-brain coverage, spanning from ventricles to SAS. Our proof-of-concept results demonstrate repeatable flow measurements and show that cardiac pulsation induces coherent CSF flow changes within the SAS. Our data also suggest that cardiac pulsation has a stronger driving effect on brain-wide CSF flow compared to respiration. This technique provides a valuable tool for investigating CSF dynamics and pathways to advance a holistic understanding of brain-wide CSF flow.

**Teaser:** A novel MR technique enables noninvasive, quantitative mapping of slow cerebrospinal fluid flow in the subarachnoid space across the human brain.

## INTRODUCTION

Cerebrospinal fluid (CSF) flow plays an important role in brain waste clearance. Early theories of CSF circulation suggested that CSF flows through the subarachnoid space (SAS) and is reabsorbed from the SAS into the venous system through arachnoid granulations (*1-3*). Recently, the glymphatic theory (*4*) has been proposed that describes an efficient pathway for waste clearance: CSF flows from the SAS into perivascular spaces (PVSs) around cerebral arteries, combining with interstitial fluid (ISF) and parenchymal solutes, and finally exiting through the venous system (*4-6*). However, our current understanding of CSF flow dynamics in the human brain is mostly limited to the ventricular system (*7-11*), such as the CSF flow in the aqueduct and fourth ventricle. Little is known about CSF flow in other regions, including the SAS and PVSs, which are critical channels for CSF circulation spanning the entire brain, but remain out of reach of current flow measurement tools. The spatial trajectory of CSF flow across the human brain has therefore not yet been measured. This poses a major barrier to understanding the CSF physiology of the human brain and its central role in waste clearance.

Another key question is what are the major driving factors of the CSF flow, as this can provide important insight into the underlying mechanisms of CSF clearance—and thereby guide the development of new therapies to enhance CSF flow and promote such clearance (*12, 13*). Both animal models and human studies (focused on CSF in the ventricles) showed that systemic physiological dynamics in the brain induced by cardiac pulsation (*8, 14-16*) and respiration (*17-19*) can modulate CSF flow. Various physiological dynamics regulate blood outflow/inflow and blood volume (*20*), which can subsequently lead to CSF inflow/outflow in ventricles to maintain constant intracranial pressure according to the Monro-Kellie doctrine (*21*). However, the effects of these systemic physiological factors, such as cardiac pulsation and respiration, on brain-wide CSF flow in the SAS remain unclear.

Magnetic Resonance Imaging (MRI) is a promising imaging tool for non-invasive flow measurement. Several MRI contrast mechanisms have been investigated to probe CSF movement, including diffusion-weighted imaging (*22-25*) and in-flow or spin labeling imaging (*11, 18*), which are magnitude-signal-based approaches, as well as phase-contrast imaging (*14, 15, 26*) which is phase-signal-based approach. Among them, phase-contrast imaging (*27*) is a quantitative approach that can directly measure the quantitative flow information including flow velocity and direction. However, conventional phase-contrast methods have limited sensitivity for detecting slow flow (e.g., < 1 mm/s). Consequently, most existing phase-contrast MRI studies focused on fast flow dynamics such as blood flow (*28-31*) and CSF flow in the aqueduct and ventricles (*9, 15, 26*), with a maximum velocity typically over 1 cm/s. Regions with slow CSF flow, such as the SAS, have not been thoroughly investigated. Indeed, the velocity range of subarachnoid CSF flow in the human brain remains unknown due to the limitations of available imaging tools.

To investigate CSF flow dynamics across the whole brain—from the ventricles to the challenging SAS—we developed a CSF flowmetry method based on phase-contrast MRI that can measure 4D CSF flow with high sensitivity, specificity, and spatiotemporal resolution. It utilizes a repurposed pulsed-gradient spin-echo (PGSE) (*32, 33*) sequence, typically used for diffusion-weighted imaging, adapted to provide high sensitivity to slow flow by achieving reduced velocity encoding (VENC) values with a long velocity-encoding time. This “snapshot” spin-echo acquisition, together with specialized phase-contrast flow data processing, provides high sampling efficiency and robustness to nuisance physiological noise.

In this proof-of-concept study, we validated the capability of our developed CSF flowmetry technique for reliable slow-flow quantification on both slow-flow phantoms and healthy volunteers. This technique successfully detected the slow CSF flow dynamics in the SAS (e.g., ∼100 μm/s), where the conventional phase-contrast method was inadequate. By applying three velocity-encoding directions (along *x, y*, and *z*), this approach provided quantitative flow velocity and direction maps with whole-brain coverage, and with good test-retest repeatability and consistency across subjects. Using this tool, we investigated the cardiac-pulsation- and respiration-coupled CSF flow dynamics in the SAS, and demonstrated that both factors can modulate CSF flow, with cardiac pulsation having a stronger driving effect than respiration.

## RESULTS

### Slow-flow-sensitized acquisition

We utilized a pulsed-gradient spin-echo (PGSE) (*32, 33*) sequence with echo planar imaging (EPI) readout (*34*) to acquire phase-contrast data (Fig. 1A). It employs a pair of motion-encoding gradients before the readout to encode the flow velocities along a given spatial direction. The use of PGSE for flow encoding allows for a relatively long encoding time (or pulse time-interval, Δ) compared to conventional flow-encoding methods, and therefore achieves low VENC values (e.g., 1.6 mm/s versus 100 mm/s), given that VENC is inversely proportional to Δ. The long encoding time also alleviates the need for increasing the amplitude (*G*) and duration (δ) of the velocity-encoding gradients to achieve the desired VENC, reducing the diffusion weighting (i.e., the effective b-value) and associated signal dephasing, which would compromise the signal-to-noise ratio (SNR) of CSF signals. The detailed relationship between VENC, b-value, Δ, G, and δ in PGSE is provided in the Methods section. To improve specificity and avoid confounding flow signals from blood, we employed long TE values (e.g., 88 ms at 7 Tesla) to obtain a strong T_2_ weighting, effectively suppressing signal from blood and tissue. Since the T_2_ value of CSF is much longer than that of blood, this minimizes the contribution of blood flow to the CSF flow quantification.

**Fig. 1.**
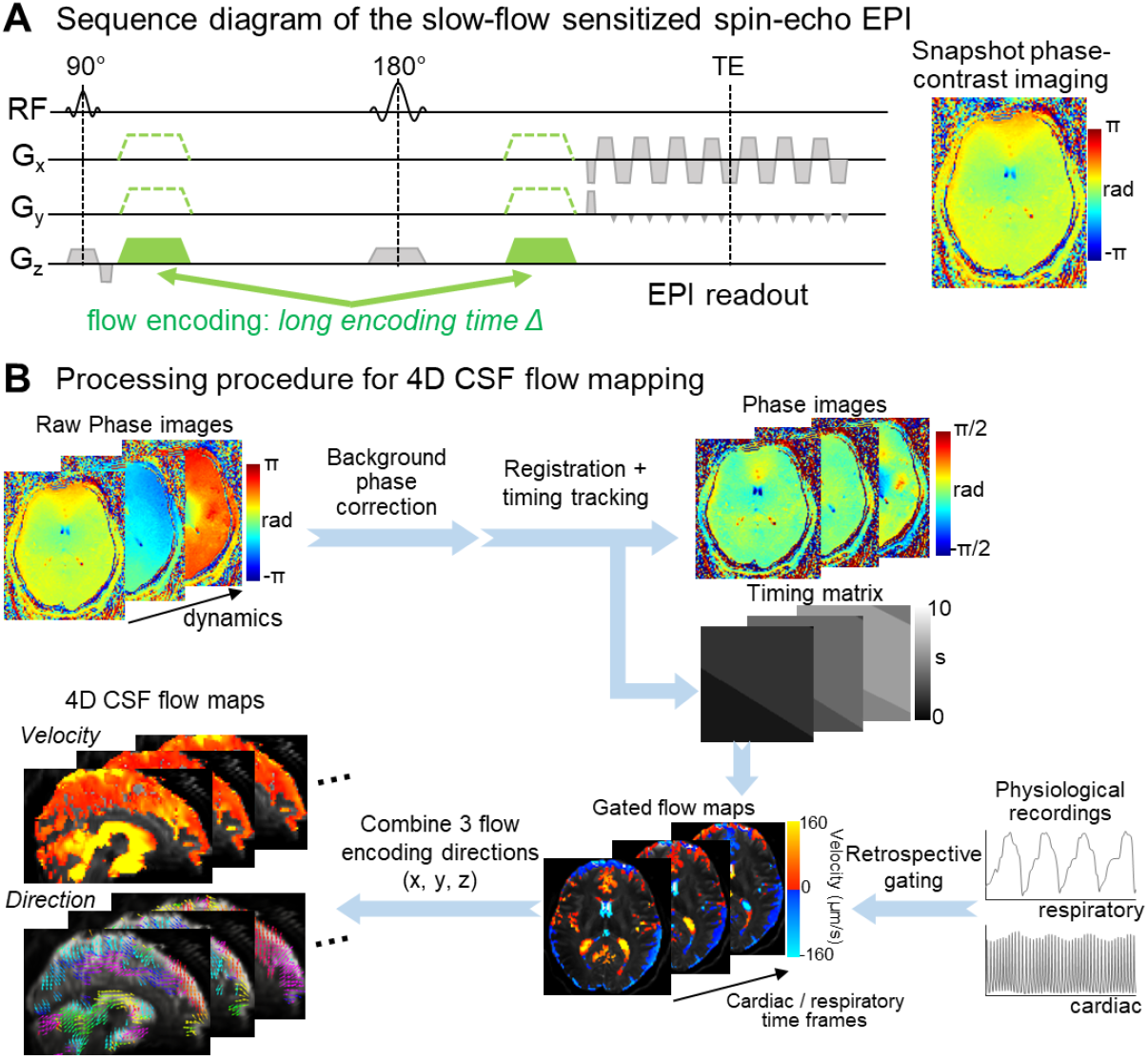
Illustration of the data acquisition and processing for CSF flowmetry. (**A**) The pulse sequence diagram of the PGSE-EPI sequence for slow-flow quantification. (**B**) The phase-contrast data processing procedure to generate 4D CSF flow velocity and direction maps from the raw phase-valued images.

In addition to improved sensitivity and specificity, PGSE-EPI provides other advantages for CSF flow mapping: (i) fast sampling provided by the EPI readout for high spatiotemporal resolution and/or large brain coverage; (ii) snapshot velocity-encoding through a single-shot acquisition to avoid shot-to-shot variations; (iii) less vulnerability to nuisance physiological noise in the image phase (e.g., caused by respiration-induced *B*_0_ field variations) provided by the spin-echo acquisition compared to conventional gradient-echo (GRE) acquisitions.

### Phase-contrast imaging data processing for 4D CSF flow mapping

We implemented a phase-contrast imaging data processing pipeline (Fig. 1B) to calculate quantitative 4D CSF flow velocity and direction maps from the raw phase-valued images. First, background-phase correction was performed by removing a low-order spatial polynomial fitted to the phase images (e.g., third order) from each timepoint independently. Background-phase correction is critical for reducing nuisance spatiotemporal variations in the phase-contrast measurements, such as induced phase changes caused by eddy currents, head motion, and residual physiological variations (e.g., respiration). Each of these nuisance phase changes is relatively spatially smooth such that they can be effectively removed by low-polynomial-order fitting, while the flow-induced phase changes of interest are spatially sparse. After background-phase removal, rigid head motion correction was performed, and, to enable proper retrospective gating, we tracked the acquisition timing of each voxel and frame based on the slice timing and estimated rigid head motion parameters. Based on the acquisition timing of each voxel and concurrent peripheral recording of systemic physiology, cardiac-or respiratory-gated phase maps can be obtained. By acquiring flow-encoding data independently along 3 spatial directions (*x, y*, and *z*), we can measure the flow speed along each direction and combine them to obtain the 4D (*x*-*y*-*z*-time) CSF flow velocity and direction.

For clarity, we define the flow-direction polarity along three spatial axes as follows: the *x*-direction represents left-right (L–R) flow, with positive values indicating flow from left to right; the *y*-direction represents posterior-anterior (P–A) flow, with positive values indicating flow from posterior to anterior; and the *z*-direction represents foot-head (F–H) flow, with positive values indicating flow from foot to head (upward).

### Custom slow-flow phantom validation

To evaluate the capability of the proposed method for accurately measuring slow flow, we first acquired data on a slow-flow phantom built around a calibrated perfusion pump (Fig. 2). We applied three different velocities of water flow (100 μm/s, 250 μm/s, 500 μm/s) through the phantom tubes (tube #1–4) arranged along the *z* direction (F–H direction), and acquired phase-contrast data using our PGSE-EPI sequence with protocol parameter values similar to those used for our *in-vivo* measurements.

**Fig. 2.**
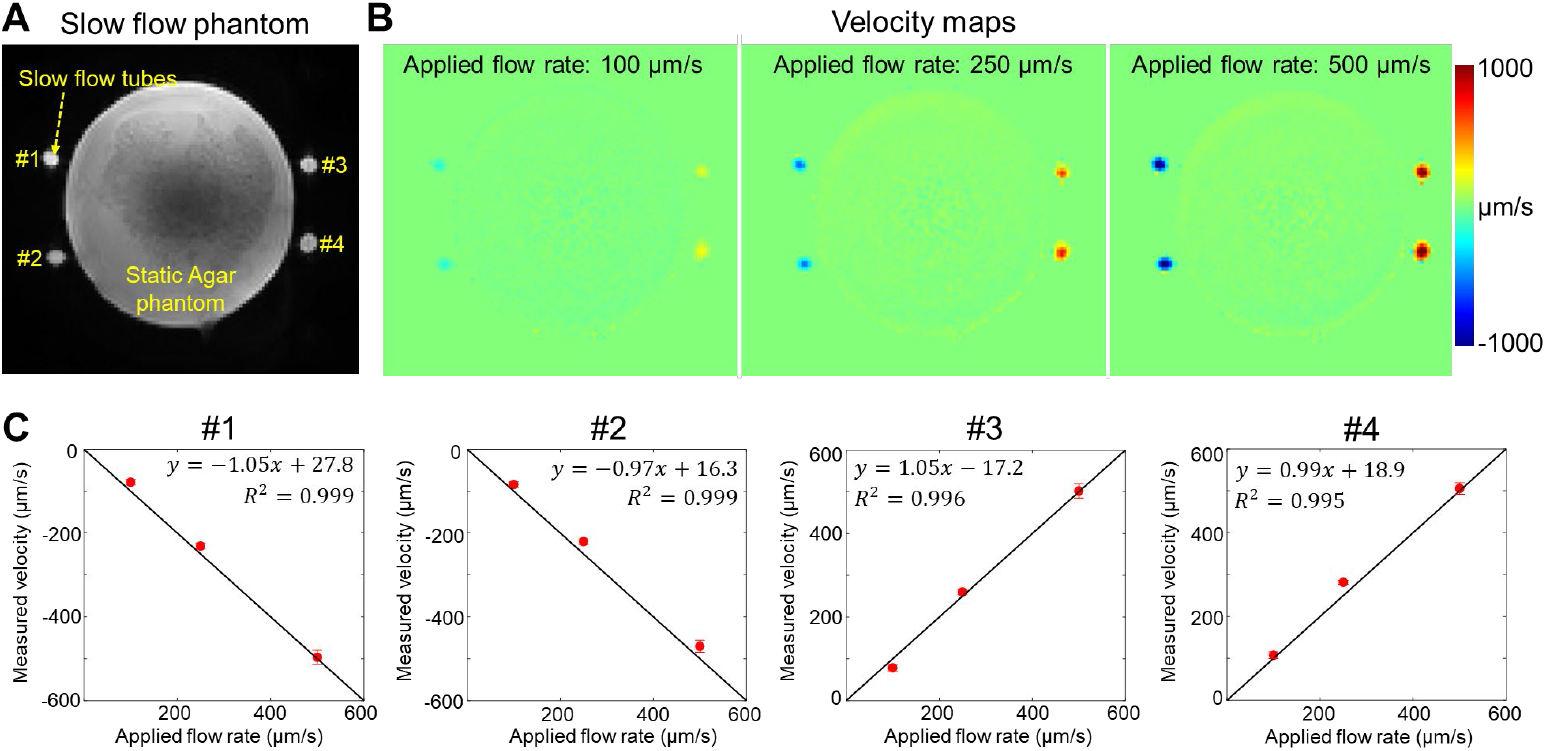
Slow-flow phantom results. (**A**) Magnitude image of the slow-flow phantom, which consists of 4 tubes (#1–4) surrounding a static agar gel phantom. The water within the tubes flows along the *z* direction, and the water in the left (#1 & #2) and right (#3 & #4) tubes flows in opposite directions. (**B**) The *z*-directed velocity maps measured at 3 different applied water flow rates (100 μm/s, 250 μm/s, and 500 μm/s). (**C**) The scatter plots of the applied water velocity versus measured velocity, along with the identity line. The dot and error bar at each data point represent the mean value and the standard deviation of the velocities across 10 timeframes within each tube’s ROI.

Figure 2B shows the measured velocity maps. The opposite signs of measured flow velocity on the left and the right sides of the phantom reflected the opposite flow direction of the water (which flows up in tubes #1&2 and down in #3&4). The increase in velocity across three measurements was also captured. Fig. 2C quantitatively assessed the accuracy of the flow measurements—the velocity plots demonstrate that the measured mean velocities agreed well with the applied velocity, with the absolute values of Pearson’s correlation coefficients (PCC) larger than 0.99 in all tubes of the phantom. The relatively small standard deviations (error bars) of the velocities across 10 timeframes suggest the stability of the measurements.

### *In-vivo* evaluation of slow CSF flow measurement

To evaluate this method in humans, high temporal resolution data were acquired with a paced breathing task (0.1 Hz, 5-s breathe in, and 5-s breathe out), because breathing tasks are known to drive CSF flow in the ventricles (*17-19*). Figure 3A shows the measured time series of velocity along the *z* direction in three representative regions-of-interests (ROIs). The time series for the ROI in the SAS (Fig. 3a) exhibited periodic dynamic changes with a maximum velocity of ∼150 μm/s. Its frequency spectrum revealed high energy at 0.1 Hz, corresponding to the frequency of the paced breathing task. As a control, the ROI in the surrounding parenchyma (Fig. 3b) did not show clear signal changes at 0.1 Hz. This suggests that the observed signal change in the SAS was unlikely to be caused by physiological noise (e.g., respiration-related *B*_0_ field changes), but rather by CSF flow changes associated with respiration. Similar respiration-associated periodic velocity changes at 0.1 Hz were also observed in the 4th ventricle (Fig. 3c) with a higher maximum velocity (∼1.5 mm/s), as expected. In addition, the CSF flow in the 4^th^ ventricle exhibited strong variations associated with cardiac pulsation at higher frequencies (> 1 Hz). The cardiac pulsation-induced flow changes appear in the time series as rapid peaks riding on the slower, periodic respiratory waveform.

**Fig. 3.**
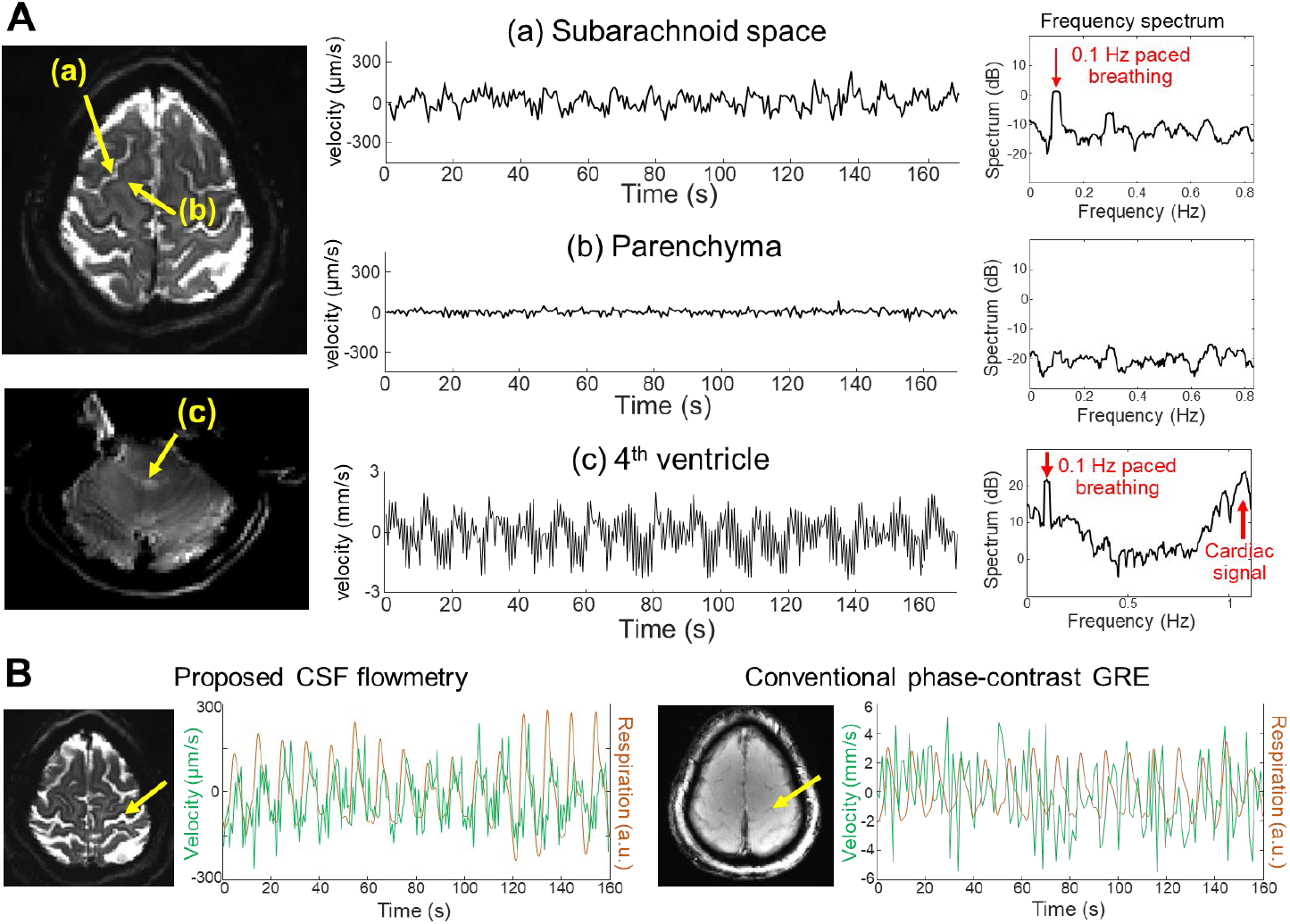
*In-vivo* high-temporal-resolution CSF flow imaging results. (**A**) Example signals measured by CSF flowmetry in the subarachnoid space (a), parenchyma (b), and 4^th^ ventricle (c) in a paced-breathing task experiment (breathing rate of 0.1 Hz), including the time-series data of the *z*-directed velocity and their corresponding frequency spectra. (**B**) Comparison of the time series (green traces) measured by either CSF flowmetry or conventional phase-contrast GRE within the subarachnoid space in the location indicated by the arrow; their synchronization (or lack thereof) with the respiration signals can be seen by inspection (orange traces).

Figure 3B compares the CSF flow dynamics in the SAS measured by the proposed CSF flowmetry with those by the conventional phase-contrast GRE sequence (with high VENC and low sensitivity to slow flow). The time series acquired by the conventional phase-contrast GRE did not agree well with the respiratory recordings and appeared as noise. In contrast, the proposed CSF flowmetry successfully detected respiration-associated flow dynamics, indicating its improved sensitivity to quantify slow flow.

All presented CSF flowmetry estimates were generated after background-phase correction. Figure S1 demonstrates that background-phase correction effectively removed the confounding physiological noise and improved the SNR of the time-series data. Before the correction, a small 0.1-Hz peak appeared in both the SAS and parenchyma (same ROIs as in Fig. 3A, a&b), likely due to the respiration-induced physiological noise (i.e., *B*_0_ field variations). After correction, the 0.1-Hz peak was diminished in parenchyma, while the spectrum of SAS showed a more distinct peak at 0.1 Hz.

### 4D CSF flow velocity and direction mapping in the SAS

We next aimed to measure the brain-wide CSF flow driven by cardiac and respiratory pulsations. We acquired whole-brain CSF flow imaging data to calculate the cardiac-and respiratory-gated CSF flow dynamics from ventricles to the brain-wide SAS. The same paced-breathing tasks were used to induce a more consistent respiratory pattern in the subjects.

Figures 4A & 4B show example maps and time-series plots of the z-directed (F–H) flow velocity across cardiac and respiratory cycles. Data from both ventricles and SAS showed strong and rapid CSF flow changes across the cardiac cycle (e.g., upward/downward flow during diastole/systole in the cisterns and ventricles around the brain stem), while relatively weaker (smaller amplitude) and slower changes were observed with the respiratory cycle. Moreover, different time delays of the velocity responses were also observed in different SAS regions (Fig. 4B). Figure 4C shows the example flow velocity maps during diastole and systole along three orthogonal flow encoding directions (L–R, P–A, and F–H). In all three flow-encoding directions, there were distinct flow direction changes (red vs. blue) between diastole and systole in both the ventricles (e.g., the atrium of the lateral ventricles) and the SAS (e.g., the anterior region of the midsagittal plane) as indicated by the white arrows.

**Fig. 4.**
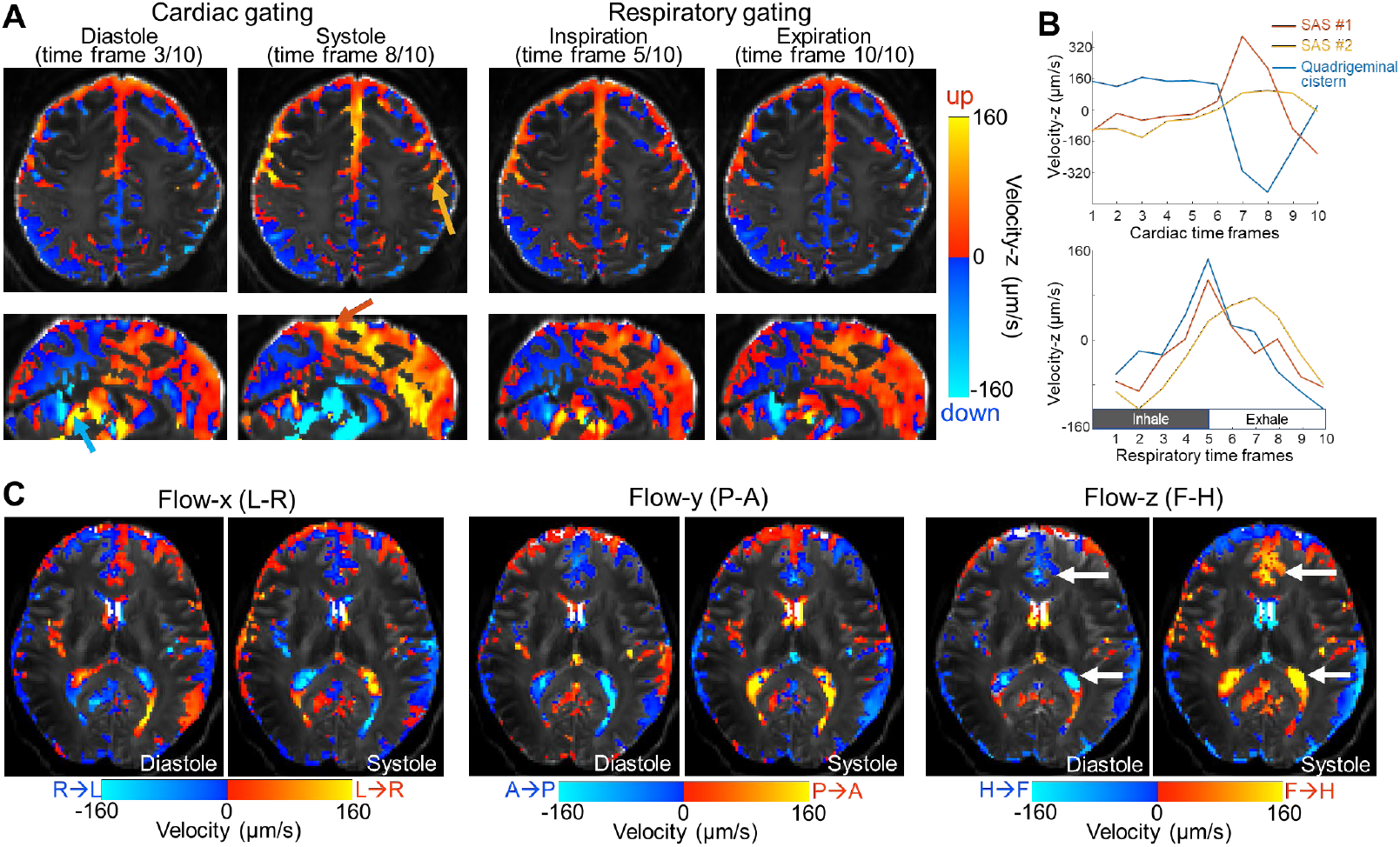
Example retrospectively cardiac-gated and respiratory-gated CSF flow results acquired using the whole-brain imaging protocol from one example subject. (**A**) *Z*-directed velocity maps at representative cardiac time frames (left panel, diastole and systole) and respiratory time frames (right panel, inspiration and expiration). Arrows in orange, red, and blue indicate three selected SAS ROIs. (**B**) Time series of the *z*-directed velocity in three selected SAS ROIs (SAS #1, SAS #2, and Quadrigeminal cistern), color-coded to match the arrows in panel A and plotted as a function of the cardiac or respiratory cycle. (**C**) Flow velocity maps measured along three different directions (*x, y*, and *z*) during diastole and systole.

Figure 5 presents the resolved velocity and flow direction changes within the cardiac cycle obtained through 3-directional velocity-encoding acquisitions. As shown in Fig. 5A, the CSF flow velocity changed across the cardiac cycle: flow velocity was relatively fast during systole (cardiac time frame 9) in both ventricles and the SAS. The color-coded vector field maps (normalized length of the 3D vector projected onto the 2D midsagittal plane) shown in Fig. 5B highlight the change of flow velocity directions (indicated by white arrows) across different time frames during the cardiac cycle. The flow velocity directions in the ventricles are consistent with previous literature (*15*): the CSF appears to flow into the brain during diastole and flows out during systole due to the outflow and inflow of the blood. There were also flow velocity direction changes detected in the SAS within the cardiac cycle between diastole and systole, and the spatial pattern of such direction changes was consistent across subjects (Fig. 5C). Animations of the color-coded vector field maps and the velocity maps of two example subjects are provided in Supplementary Movie S1 and S2 to depict the dynamic changes of the CSF flow across the cardiac cycle.

**Fig. 5.**
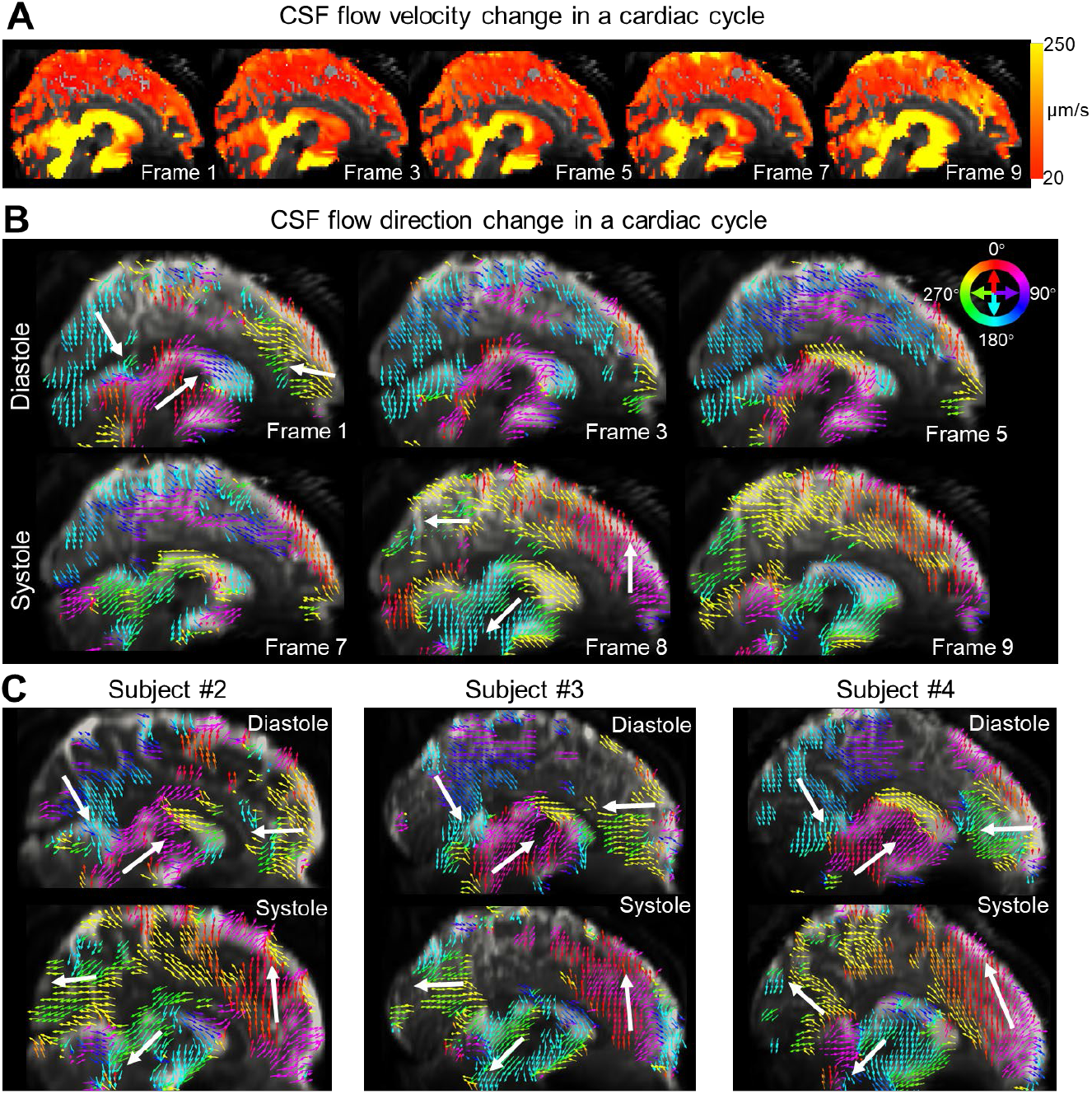
Examples of 4D CSF flow velocity and direction maps displayed in the midsagittal plane. (**A**) CSF flow velocity maps at different time frames of the cardiac cycle. (**B**) Velocity vector field maps at different cardiac time frames. The length of the 3D vector is normalized and projected onto the 2D midsagittal plane. The angle and color of the vectors both represent flow direction. A threshold of 20 μm/s was applied to exclude slower flow. (**C**) Velocity vector field maps from three additional volunteers during diastole and systole of the cardiac cycle. The white arrows highlight the consistent flow velocity direction changes between diastole and systole across subjects.

### Scan-rescan repeatability evaluation

To evaluate the repeatability of the proposed CSF flowmetry, we assessed scan-rescan reproducibility on 6 healthy volunteers (23 to 46 years old) using our whole-brain PGSE-EPI acquisition. The 3-directional velocity encoding was acquired with an acquisition time of ∼13 minutes for each scan, and 2 scans were performed on each subject. Retrospective cardiac gating was performed to estimate velocity range (*V*_range_, defined as the difference between maximum velocity and minimum velocity) throughout the cardiac cycle along 3 directions. This velocity range served as a quantitative metric to evaluate the repeatability of flow velocity measurements. Figure 6A shows example z-directed *V*_range_ maps of a pair of test-retest scans acquired on a single subject, as well as their difference maps.

**Fig. 6.**
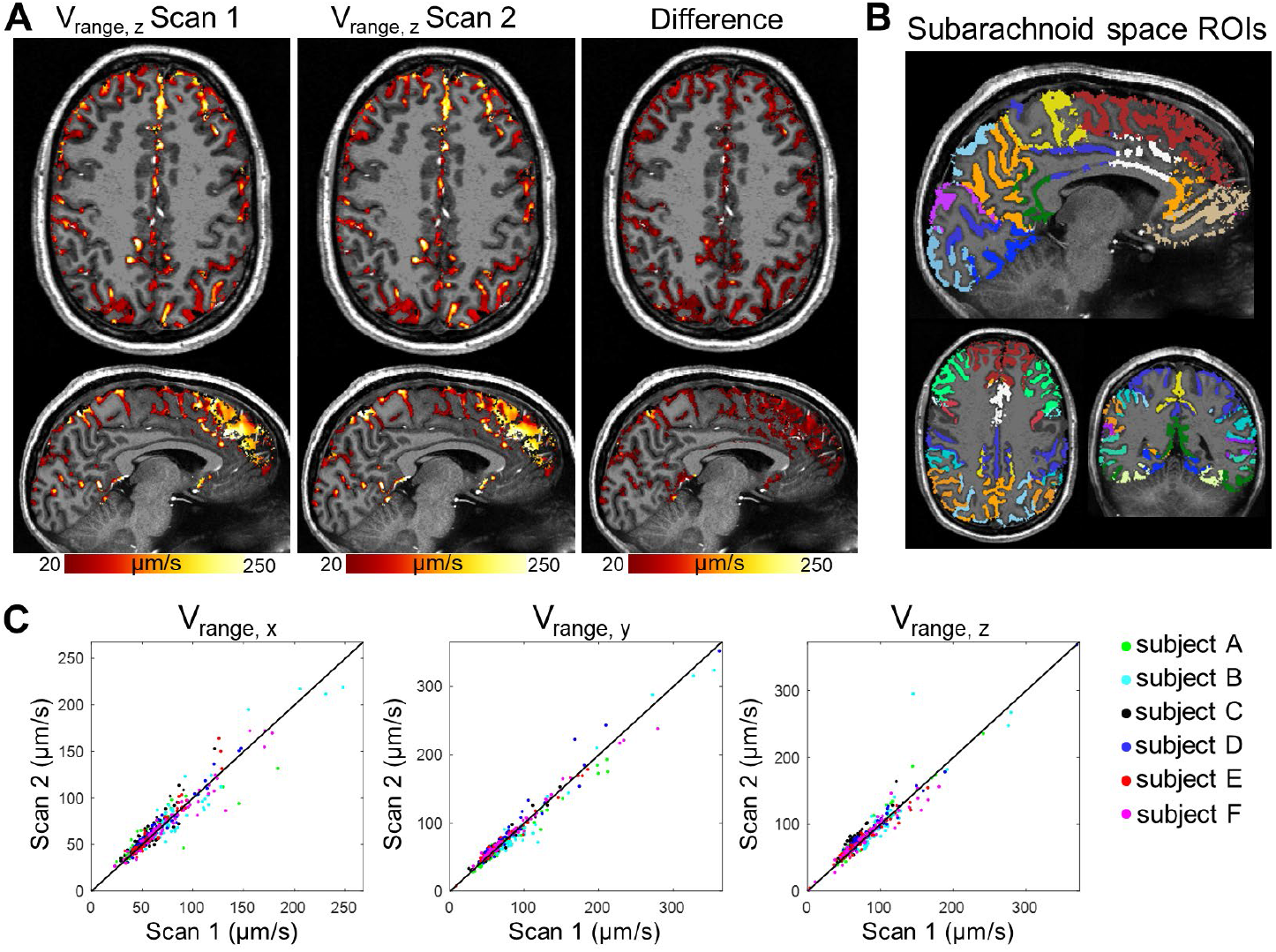
Test-retest repeatability results of SAS CSF flow. (**A**) The *V*_range_ maps of z-directed flow velocity from the two repeated scans (i.e., one scan-rescan pair) and the corresponding difference maps on an example volunteer. *V*_range_ quantifies the range of velocity during the cardiac cycle, defined as maximum velocity minus minimum velocity across the cycle. (**B**) The segmented SAS ROIs used for repeatability analyses. (**C**) Scatter plots of the test-retest *V*_range_ values in the 70 ROIs measured from 6 healthy subjects, shown along with the identity line for reference.

We performed ROI-based test-retest analyses of CSF flow in the SAS across different brain regions using ROIs obtained from FreeSurfer. Figure 6B shows the segmented SAS ROIs of an example subject. The scan-rescan data exhibited strong correlations (high PCC values) for *V*_range_ along all three velocity encoding directions as shown in Fig. 6C: PCC = 0.96 for *V*_range_ along *x* (*p* < 0.0001), PCC = 0.98 for *V*_range_ along *y* (*p* < 0.0001), and PCC = 0.95 for *V*_range_, along *z* (*p* < 0.0001). In addition, we also analyzed the repeatability of the standard deviation of the velocities measured across the cardiac cycle (*V*_SD_) as another metric, reflecting the degree of temporal variation in flow dynamics. Similarly, the scan-rescan *V*_SD_ along all 3 velocity encoding directions showed strong correlations as shown in Fig. S2, indicating good repeatability of the flow measurements in SAS.

### The effects of cardiac pulsation and respiration

We analyzed the cardiac-gated and respiratory-gated flow velocity metrics (*V*_range_ and *V*_SD_) using the same SAS ROIs in healthy volunteers (*N* = 6) to investigate the effects of cardiac pulsation and respiration on modulating brain-wide CSF flow dynamics.

Figures 7A & B present *V*_range_ and *V*_SD_ calculated along the 3 velocity-encoding directions from cardiac-gated flow measurements, respiratory-gated flow measurements acquired during a 0.1-Hz paced breathing task, and respiratory-gated flow measurements acquired during free breathing. The cardiac-gated flow velocity showed significantly higher overall *V*_range_ and *V*_SD_ values than respiration-gated flow velocity (*p* < 0.0001) across all 3 velocity-encoding directions. Paced breathing produced a slightly higher *V*_range_ compared to free breathing (*p* = 0.0011). Among the directions, z-directed flow (F–H) exhibited the highest *V*_range_ and *V*_SD_ values, while *x*-directed flow (L–R) showed the lowest. Example *V*_range_ maps from two representative subjects are presented in Fig. 7C, highlighting the higher values in cardiac-gated maps compared to respiration-gated maps. The maps also showed the relatively large *V*_range_ along the *z* direction and the low *V*_range_ along the *x* direction. These results suggest that cardiac pulsation has a stronger driving effect than respiration on brain-wide CSF flow dynamics in the SAS.

**Fig. 7.**
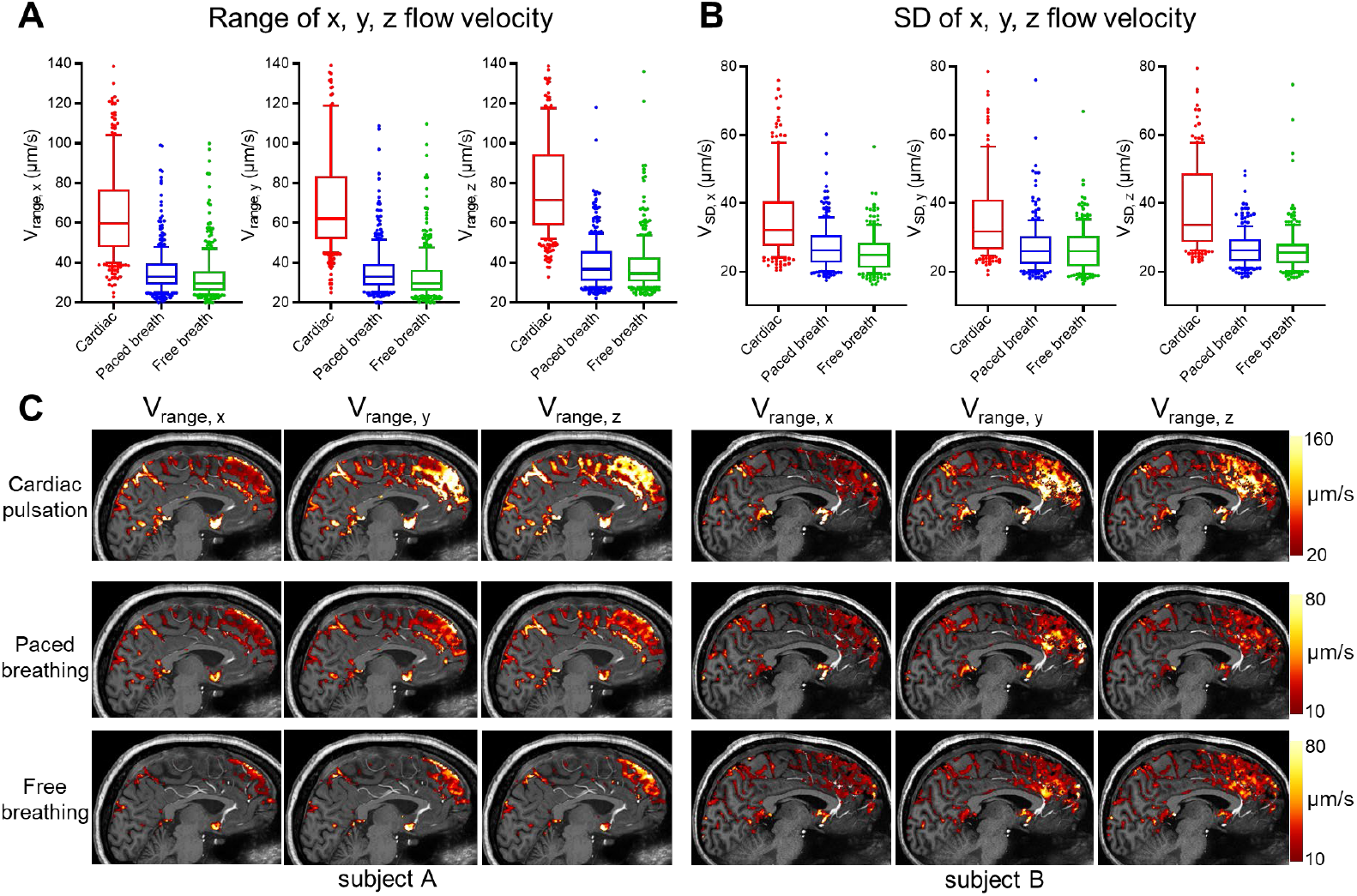
Comparison of the effects of cardiac pulsation and respiration on SAS CSF flow. (**A**) Box- and-whisker plots (N=6) showing median, interquartile range, and 90% range of the *V*_range_ values (across ROIs and volunteers) grouped by different modulating factors, including cardiac pulsation, respiration with paced breathing, and respiration with free breathing. Plots for *V*_range_ along the 3 velocity-encoding directions (*x, y*, and *z*) are presented. (**B**) Similar box-and-whisker plots of the *V*_SD_ values along 3 velocity-encoding directions, where *V*_SD_ represents the standard deviation of the velocity values measured across either cardiac or respiratory cycle. (**C**) *V*_range_ maps obtained through cardiac-gating (cardiac pulsation) and respiratory-gating (paced and free breathing) along the 3 velocity-encoding directions from two example subjects.

## DISCUSSION

In this study, we developed a CSF flowmetry technique to map CSF flow dynamics across the whole brain from ventricles to SAS. This technique achieves high sensitivity and efficiency for slow-flow mapping through slow-flow-sensitized PGSE-EPI acquisition and dedicated phase-contrast flow velocity data processing. Both the phantom and *in-vivo* experiments demonstrated sensitivity for measuring slow flow (e.g., ∼100 μm/s), enabling the investigation of brain-wide CSF flow dynamics in SAS. It provides quantitative measurements of flow velocity and direction with high sampling efficiency. These proof-of-concept results demonstrated that the proposed CSF flowmetry was able to map 4D CSF flow dynamics and their changes across cardiac and respiratory cycles. Additionally, our results exhibited good repeatability with high PCC values in preliminary scan-rescan experiments on healthy volunteers.

The high sensitivity for mapping slow CSF flow relies on both data acquisition and processing. On the acquisition side, PGSE-EPI provides the flexibility to use a long velocity-encoding interval to achieve low VENC values without necessitating strong diffusion gradients that could diminish the CSF signal. In addition, PGSE-EPI offers several advantages over conventional phase-contrast MRI for the purpose of reducing physiological noise, including the use of single-shot EPI to avoid shot-to-shot signal/phase variations, and the use of spin-echo instead of gradient-echo acquisition to improve robustness against *B*_0_ field fluctuations. These features are critical for reliable phase-contrast measurements of slow flow, as the phase changes caused by slow CSF flow are much smaller than typical fast blood or ventricular flow, making the signals more vulnerable to physiological noise induced by respiration or motion. On the processing side, the background-phase correction further reduces physiological noise contributions in the phase measurements, as illustrated in Fig. S1.

CSF flow dynamics are complex and can be modulated by multiple driving factors. Previous studies suggested that CSF flow measured in human ventricles or the aqueduct can be driven by cardiac pulsation (*8, 14, 15*), respiration (*17-19*), and neuronal activity (*11, 35, 36*). With our CSF flowmetry technique, we can investigate the relationship between brain-wide CSF flow dynamics and these potential driving factors (*8, 15*), and identify specific flow dynamics and pathways associated with different physiological mechanisms. In this proof-of-concept study, we investigated the effects of cardiac pulsation and respiration on brain-wide CSF flow using retrospectively-gated flow metrics. These preliminary data suggest that both cardiac pulsation and respiration modulate CSF flow in the SAS, and that cardiac pulsation has a stronger driving effect than respiration. Paced-breathing tasks also enhanced flow dynamics more than free breathing, although the effect was modest. In addition, we observed distinct changes in flow velocity direction across the cardiac cycle (Fig. 5). Future work will further investigate CSF flow patterns and pathways in the SAS across different brain regions.

Another potential application of CSF flowmetry is in studying abnormalities of CSF flow in neurodegenerative disease. Recent animal studies suggest that a normal-functioning CSF system is crucial for maintaining brain health across the lifespan (*37, 38*), and disruptions in CSF flow may contribute to the development of dementia and other disorders (*4, 39-41*). The proposed CSF flowmetry offers an efficient tool for noninvasively imaging CSF in the human brain, providing quantitative 4D CSF flow measurements with whole-brain coverage in a clinically acceptable timeframe (e.g., 3-directional flow protocol in 13 minutes). The high efficiency and quantitative measurements make it a valuable tool for investigating CSF flow dynamics in clinical settings, to identify pathological flow dynamics or to track treatment response.

As imaging brain-wide CSF flow is a new direction, there are many avenues for improvement, from data acquisition to post-processing. For example, in addition to mapping CSF flow in SAS as shown in this study, the developed CSF flowmetry approach could be applied to map CSF flow in PVSs. To achieve this, higher spatial resolution would be required to image the small PVSs and reduce the partial volume effects from surrounding tissues. Advanced PGSE-based acquisition techniques that can address distortion and blurring artifacts in EPI at high spatial resolutions (*42, 43*) and enhance SNR efficiency (*44, 45*) present promising avenues for increasing the spatial resolution and specificity of CSF flowmetry for PVS imaging. For post-processing, improved segmentation tools tailored for CSF spaces are needed for more accurate analysis of the SAS and PVSs. Better visualization techniques to capture the sparse, whole-brain CSF flow in the SAS and PVSs and their interconnectivity could also facilitate the interpretation and quantitative analysis of these rich 4D data. Finally, net flow information can be derived from the quantitative flow velocities, which may offer insights into CSF flow pathways and waste transport. Future work will focus on these directions to further investigate the brain-wide CSF flow patterns and circulation pathways.

In conclusion, we presented a CSF flowmetry technique that enables quantitative 4D CSF flow imaging with high sensitivity to slow flow and whole-brain coverage. This technique successfully measures the slow CSF flow dynamics in the SAS across the brain, and their velocity and direction changes across the cardiac and respiratory cycles. This technology provides a valuable tool for investigating brain-wide CSF flow dynamics and patterns in humans to obtain insights into CSF physiology and dysfunction in health and disease.

## MATERIALS AND METHODS

### VENC value and diffusion weighting in PGSE

VENC is a key parameter in phase-contrast MRI that represents the maximum velocity the sequence can measure without introducing phase wrapping of the phase signals. The VENC value of the phase-contrast signals in the PGSE sequence is inversely proportional to both Δ and *G* as:

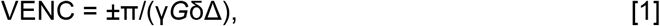

where “γ” is the gyromagnetic ratio, “G” is the amplitude of velocity-encoding gradients, δ is the gradient duration of each velocity encoding gradient, and Δ is the gradient pulse interval. A lower VENC value can improve the sensitivity to slow flow by generating stronger flow-induced phase changes. To reduce the VENC, we can increase Δ, *G*, or δ in the sequence design. However, increasing these parameters can result in stronger diffusion weighting, reflected by a higher b-value, which diminishes the CSF signal and reduces the SNR. The b-value of the PGSE is defined as:

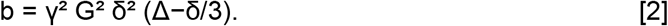

This shows that the b-value will increase quadratically with the increase of *G* and δ, but only linearly with Δ. Therefore, compared to increasing *G* and δ, increasing the velocity-encoding interval Δ is a more effective way to reduce VENC value and increase sensitivity to slow flow, while minimizing signal dephasing caused by diffusion weighting. The proposed PGSE-based phase-contrast acquisition allows for a longer velocity-encoding interval compared to conventional phase-contrast MRI, enabling low-VENC slow flow mapping.

### Experimental design and data acquisition

All volunteers provided written informed consent prior to scanning using an approved IRB protocol, following all policies of our institution’s Human Subjects Research Committee. Subjects were healthy adult volunteers (*N*=8, 22 to 46 years old, 5 female, 3 male). All data in this study were acquired on a whole-body human 7 Tesla scanner (MAGNETOM Terra, Siemens Healthineers, Erlangen, Germany). Experiments were conducted in five sessions using an in-house-built 64-channel head array coil (*46*) and in three sessions using a vendor-supplied 32-channel head array coil (Nova Medical, Wilmington, MA). Physiological signals were acquired concurrently during all scan sessions using external sensors: cardiac signals were measured with a piezoelectric device on the fingertip, and respiratory signals were captured with respiratory bellows positioned around the chest. The acquisition parameters for the MRI protocols are described below.

#### Slow-flow phantom scans

Flow quantification validation was conducted using a custom-built slow-flow phantom based on a calibrated perfusion pump feeding tap water through acrylic tubes oriented along the *z*-direction using three constant flow velocity rates (100 μm/s, 250 μm/s, and 500 μm/s). PGSE-EPI data were acquired with the following parameter values: 1.2 × 1.2 × 2 mm^3^ voxels, FOV = 192 × 192 × 64 mm^3^, VENC = 1.6 mm/s, TE/TR = 85/2500 ms, multiband factor = 2, in-plane acceleration factor (*R*_in-plane_) = 3. Ten dynamics were acquired with *z*-directed velocity encoding, and were averaged to calculate velocity.

#### High-temporal-resolution *in-vivo* scans

High-temporal-resolution data were acquired in the 4^th^ ventricle and the SAS during a paced-breathing task (frequency = 0.1 Hz, 5 s inhale, and 5 s exhale) from 2 healthy subjects. Instructions on when to breathe in and out were presented visually throughout the scan. These scans were performed with short TR values (450 ms for the 4^th^ ventricle scan, 600 ms for the SAS scan) and fewer slices to investigate velocity dynamics and their frequency spectrum at high temporal resolutions. The other acquisition parameter values were: resolution = 1.6 × 1.6 × 5 mm^3^, FOV (in-plane) = 210 × 210 mm^2^, *R*_in-plane_ = 3, no multiband, 4 slices with VENC = 4.8 mm/s and TE = 50 ms for the 4^th^ ventricle scan, and 5 slices with VENC = 1.6 mm/s and TE = 81 ms for the SAS scan. Three minutes of data were acquired for each protocol. The first 5 dynamics of the scan were acquired without velocity encoding, followed by data with z-directed velocity encoding. For comparison, conventional phase-contrast GRE data were acquired in the 4^th^ ventricle and SAS at the same resolution and location with the following parameter values: number of slices = 1, volume TR (temporal resolution) = 2.57 s, TR/TE = 47.70/5.92 ms, *R*_in-plane_ = 2, VENC = 10 cm/s, *z*-directed velocity encoding.

#### 4D whole-brain CSF flow scans

PGSE-EPI data with 3-directional velocity-encoding and whole-brain coverage were acquired to investigate cardiac-gated or respiratory-gated CSF flow through retrospective gating. Data were collected with and without paced-breathing tasks (i.e., paced breathing and free breathing) from 6 healthy volunteers (23 to 46 years old) with 2 repetitions for scan-rescan repeatability evaluation. The second scan (i.e., a repetition) was acquired ∼30 minutes after the first scan within the same session. The acquisition parameter values were: resolution = 1.6 × 1.6 × 3.2 mm^3^, FOV = 211 × 211 × 115.2 mm^3^, VENC = 1.6 mm/s, TE/TR = 81/2100 ms, multiband factor = 2, *R*_in-plane_ = 2. A total of 120 dynamics were acquired for each velocity-encoding direction, including 5 volumes without velocity-encoding and 115 volumes with velocity-encoding. The acquisition time for each direction was ∼4 min 20 s, and the total scan time for 3 directions was ∼13 minutes. In addition, *B*_0_ field maps were acquired using a standard dual-echo GRE sequence with matched spatial resolution to enable geometric distortion correction of the EPI data. Anatomical reference images used for brain segmentation were obtained using a standard T_1_-weighted MPRAGE at 0.75-mm isotropic resolution.

### Data processing and analysis

The CSF flow data processing pipeline is summarized in Fig. 1B. Initially, the ASPIRE coil combination algorithm (*47*) was used to obtain raw phase images, to avoid phase singularities and improve the quality of the phase signals. For background-phase correction, we utilized a third-order spatial polynomial fitting to estimate the background phase, based on the assumption that both eddy current-induced phase and physiological noise or motion-induced background phase changes are spatially smooth, while flow-related phase signals are spatially sparse and change abruptly in the image. To minimize bias, voxels that are likely to be CSF were excluded from the fitting by masking out high-intensity voxels, motivated by the fact that CSF has higher intensities in T_2_-weighted PGSE images. The background-phase correction was performed on each slice and frame independently. Subsequently, motion correction was performed using AFNI (*48, 49*). Motion parameters were estimated from the magnitude-valued images and applied to the phase-valued images. To ensure accurate timing for retrospective gating, the ‘nearest neighbor’ interpolation method was used, and the acquisition time of each voxel of each time frame was recorded to track the true acquisition timing after data resampling applied during motion correction. Phase values were converted to velocity V = ϕ/π × VENC, where ϕ is the phase value. Using the recorded physiological signals and acquisition timings, cardiac-gated or respiratory-gated flow velocity maps were calculated by binning the flow signals into different relative times within the cardiac or respiratory cycle; ten such bins were used for retrospective gating. Finally, by acquiring 3-directional flow velocity data, 3-dimensional velocity vector fields were generated, from which velocity magnitude and direction of the flow were calculated. A CSF mask, generated automatically by the FAST tool of the FSL software package (*50, 51*) was applied to the calculated CSF flow maps.

For test-retest repeatability evaluation, we further processed the whole-brain PGSE-EPI data with geometric distortion correction and registered them to the MRPAGE T_1_-weighted volume. Distortion correction was performed by FUGUE from FSL (*51*) with the field map estimated from the dual-echo GRE data. The MPRAGE data were segmented with FreeSurfer (*52-54*) to generate the SAS ROIs. The ‘DKTatlas’ was used to parcellate the cerebral cortex, and the segmented parcels were extended outwards from cortical gray matter to the CSF space to obtain the SAS masks. This resulted in 70 ROIs (example shown in Fig. 6B), which were used to calculate ROI-based flow velocity metrics (e.g., *V*_range_ or *V*_SD_ within each ROI) in the repeatability analysis (Fig. 6) and in the investigation of the effects of cardiac pulsation and respiration (Fig. 7). For statistical analysis, Pearson’s correlation coefficients were calculated to evaluate repeatability, and one-tailed t-tests were used for comparing the cardiac-gated and respiratory-gated (free-breathing and paced-breathing) flow metrics.

## ACKNOWLEDGMENTS

We would like to thank Kyle Droppa and Estee Perelgut for their help with subject recruitment and MRI scanning support. This work was supported by the National Institutes of Health (grants K99-AG083056, U24-NS129893, R01-EB036507, R01-AT011429, U19-NS128613, R01-AG070135, R01-EB019437, P41-EB030006), and the Athinoula A. Martinos Center for Biomedical Imaging; and was made possible by the resources provided by NIH Shared Instrumentation Grant S10-OD023637.

## Author contributions

Conceptualization: ZD, FW, JRP; Methodology: ZD, FW, JRP; Investigation: ZD, FW, JRP, AKS; Processing and optimization: ZD, FW, SDR, KE, BB; Supervision: ZD, FW, JRP, LDL, LLW, BRR; Writing—original draft: ZD, FW; Writing—review & editing: ZD, FW, JRP, LDL, SDR, LLW, BRR

## Data and materials availability

All data needed to evaluate the conclusions of this study are present in the main text and the supplementary materials.

## SUPPLEMENTARY MATERIALS

### Supplementary Figures S1–S2

**Fig. S1. Time-series data and corresponding frequency spectra of the signals measured by CSF flowmetry before and after background-phase correction**. The same data from Fig. 3 and the same ROIs in SAS (top) and parenchyma (bottom) are plotted. Before correction (left panel), the time series data from the SAS and the parenchyma ROIs are noisy, and their frequency spectra have a small peak at 0.1 Hz (red arrows). After correction (right panel), the time series are less noisy, with a more distinct 0.1-Hz peak in SAS but a diminished peak in parenchyma.

**Fig. S2. Scatter plots of the test-retest *V***_**SD**_ **values measured from *N*=6 healthy subjects, shown along with the identity line for reference**. *V*_SD_: standard deviation of the velocity within the cardiac cycle. The test-retest *V*_SD_ plots along three velocity-encoding directions are shown. PCC = 0.95 for *V*_SD_ along *x* (*p* < 0.0001), PCC = 0.98 for *V*_SD_ along *y* (*p* < 0.0001), and PCC = 0.96 for *V*_SD_ along *z* (*p* < 0.0001).

### Supplementary Movie S1–S2

**Movie. S1**. The dynamic changes of the CSF flow direction and velocity across the cardiac cycle shown by color-coded vector field maps and the velocity magnitude maps of a representative subject (Subject #1).

**Movie. S2**. The dynamic changes of the CSF flow direction and velocity across the cardiac cycle shown by color-coded vector field maps and the velocity magnitude maps of a representative subject (Subject #2).

