## Supplementary materials for "Quantifying brain-wide cerebrospinal fluid flow dynamics using slow-flow-sensitized phase-contrast MRI"

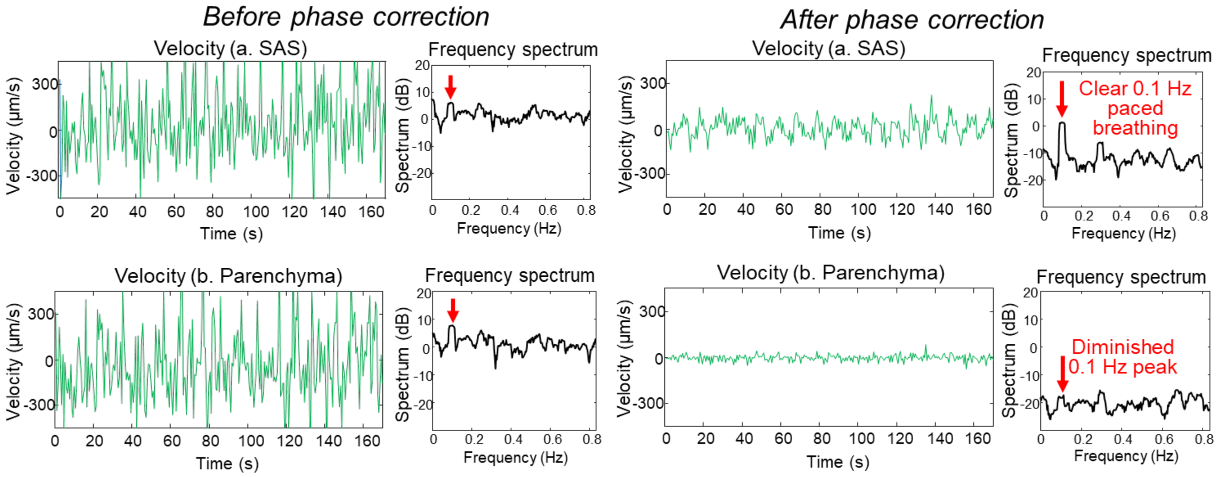


**Fig. S1. Time-series data and corresponding frequency spectra of the signals measured by CSF flowmetry before and after background-phase correction.** The same data from Fig. 3 and the same ROIs in SAS (top) and parenchyma (bottom) are plotted. Before correction (left panel), the time series data from the SAS and the parenchyma ROIs are noisy, and their frequency spectra have a small peak at 0.1 Hz (red arrows). After correction (right panel), the time series are less noisy, with a more distinct 0.1-Hz peak in SAS but a diminished peak in parenchyma.

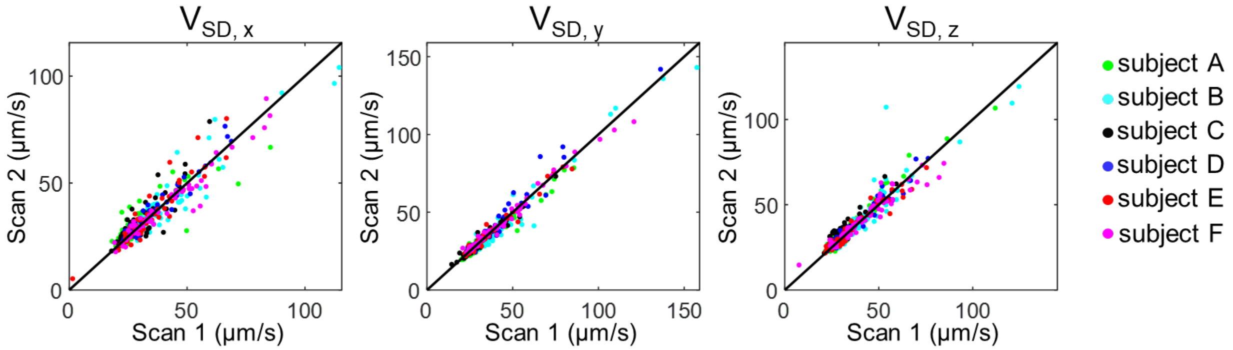


**Fig. S2. Scatter plots of the test-retest $V_{SD}$ values measured from $N$=6 healthy subjects, shown along with the identity line for reference.** $V_{SD}$: standard deviation of the velocity within the cardiac cycle. The test-retest $V_{SD}$ plots along three velocity-encoding directions are shown. PCC = 0.95 for $V_{SD}$ along $x$ ($p < 0.0001$), PCC = 0.98 for $V_{SD}$ along $y$ ($p < 0.0001$), and PCC = 0.96 for $V_{SD}$ along $z$ ($p < 0.0001$).

**Movie. S1. The dynamic changes of the CSF flow direction and velocity across the cardiac cycle shown by color-coded vector field maps and the velocity magnitude maps of a representative subject (Subject #1).**

**Movie. S2. The dynamic changes of the CSF flow direction and velocity across the cardiac cycle shown by color-coded vector field maps and the velocity magnitude maps of a representative subject (Subject #2).**
